# Power Pixels: a turnkey pipeline for processing of Neuropixel recordings

**DOI:** 10.1101/2025.06.27.661890

**Authors:** Guido T. Meijer, Francesco P. Battaglia

## Abstract

There are many open-source tools available for the processing of neuronal data acquired using Neuropixels probes. Each of these tools, focuses on a part of the process from raw data to single neuron activity. For example, SpikeInterface is an incredibly useful Python module for pre-processing and spike sorting of individual recordings. However, there are more steps in between raw data and spikes, such as synchronization of spike times between probes and histological reconstruction of probe insertions. Therefore, we developed Power Pixels, combining the functionality of several packages into one integrated pipeline, which may be run in any lab workflow. It includes pre-processing, spike sorting, neuron-level quality control metrics, synchronization between multiple probes, compression of raw data, and ephys-to-histology alignment. Integrating all these steps into one pipeline greatly simplifies Neuropixels data processing, especially for novel users who might struggle to find their way around all the available code and tools.

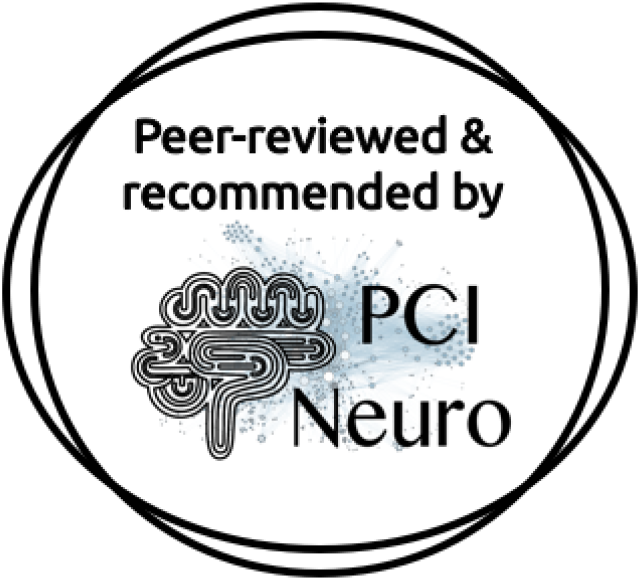

## Introduction

The processing of electrophysiological recordings, from raw data to spike-sorted neurons in defined brain regions, is a very labor-intensive task for many researchers. This in particular with the advent of high-density silicon probe, such as Neuropixels (Jun et al., 2017). Large collaborations, like the International Brain Laboratory (IBL), have developed fully integrated end-to-end pipelines, this takes the workload away from the researchers and allows them to spend more time doing experiments, analysis and writing. These pipelines, however, are embedded in the database architecture used by the collaboration and do not run out-of-the-box on recordings made by external researchers. (International Brain Laboratory et al., 2022). Even when the code base is open-source, it’s challenging for external researchers to apply the code to their data because the code is written for large-scale deployment instead of running a single local recording session. Furthermore, collaborations might have access to advanced equipment which is typically not available to individual labs.

Advances are made to streamline and standardize processing and spike sorting of electrophysiological recordings, most notably by SpikeInterface (Buccino et al., 2020). SpikeInterface is a powerful tool which is aimed at the first stages of processing: preprocessing, spike sorting, and curation of the spike sorting output. To get a full end-to-end pipeline, however, several additional steps are required, namely: synchronization between multiple probes, histological reconstruction of probe tracts, and alignment of histology to electrophysiological markers. We present here the Power Pixels pipeline, covering all these steps and additionally calculates several different neuron-level Quality Control (QC) metrics which can help, or even completely replace, manual curation of spike sorting output. In short, the Power Pixels pipeline combines SpikeInterface, the IBL pipeline, and probe tracing software into one end-to-end pipeline from raw recorded data to spike sorted neurons in defined anatomical regions in the brain.

## Methods

The Power Pixels pipeline combines processing steps from SpikeInterface, elements from the IBL pipeline, and a standalone MATLAB package into one end-to-end pipeline (Figure 1). The pipeline takes raw data electrophysiological time series as input and generates curated spike-sorted neurons, along with the anatomical location of the recorded neurons as output. The backbone of the pipeline is the *powerpixels* Python class that contains several functions related to the different steps of the pipeline. To establish in which brain region each recording channel ended up, histological reconstruction of the probe tracts is done using the MATLAB package AP_Histology or Universal Probe Finder (Montijn and Heimel, 2022). Finally, the electrophysiological signatures along the probe are aligned with the inferred brain regions from the probe tract using the [Ephys-Histology Alignment GUI](https://github.com/int-brain-lab/iblapps/wiki) developed by the IBL.

**Figure 1.**
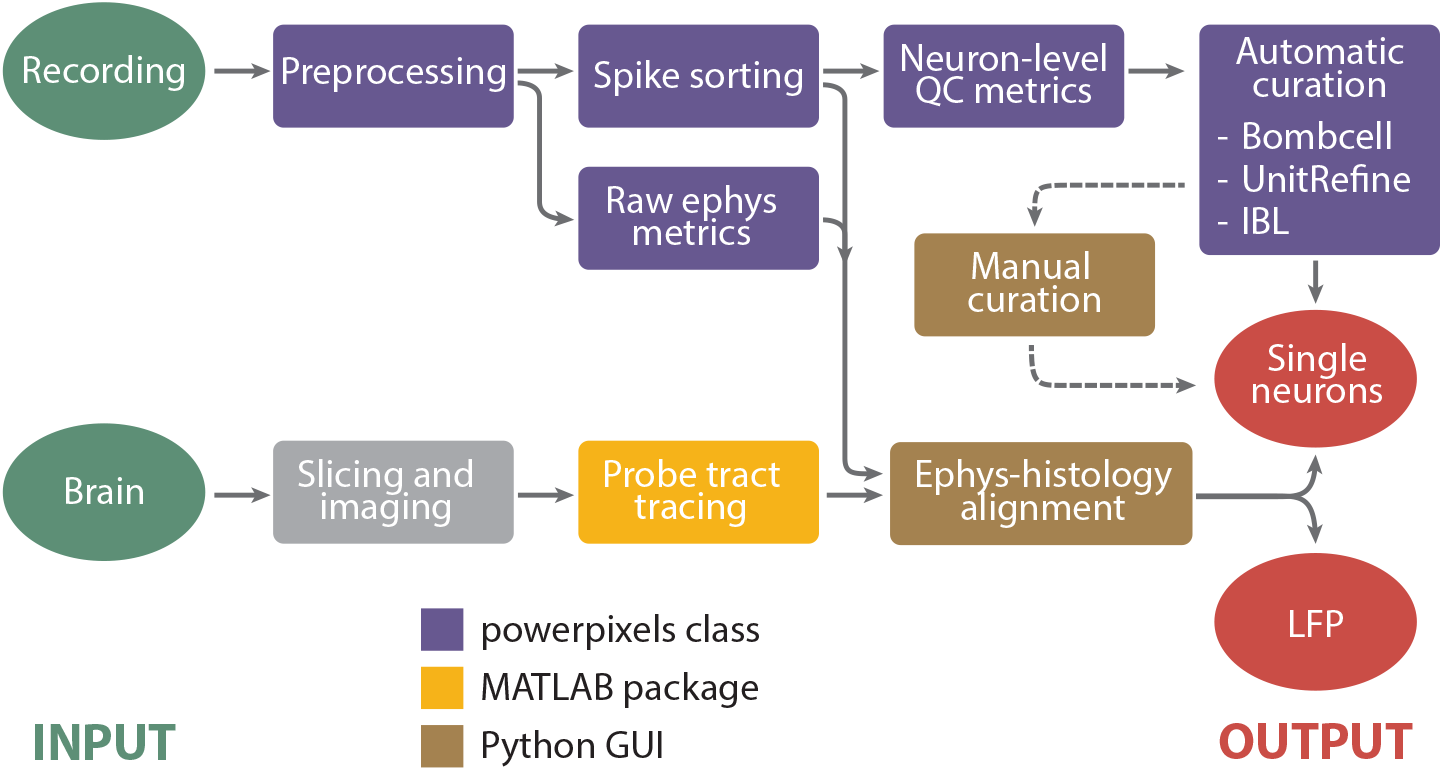
Process diagram of the pipeline. The inputs are the raw recording and the perfused brain and the outputs are single neurons and LFP power, in defined brain regions.

### Supported data formats and hardware

The pipeline supports recordings done with Neuropixels 1.0 and 2.0 using SpikeGLX or OpenEphys on a National Instruments acquisition system. Accepted raw data formats are therefore .bin and .dat files, from SpikeGLX and OpenEphys respectively. Also the compressed raw data formats .cbin and .zarr are accepted, these are decompressed before the pipeline is run because some steps require uncompressed raw binary data (e.g. Bombcell). Other Neuropixels probes, like the 1.0 NHP, 2.0 Quad Base, or Ultra, will most likely be accepted by the pipeline but have not been tested. Non-Neuropixel probes, like the SiNAPS from NeuroNexus (Angotzi et al., 2019), are less likely to work out-of-the-box. The same goes for other acquisition devices, like the OneBox, they have not been tested but might work with some manual tweaking.

### Preprocessing

Long-shanked, high-density silicon probes, such as Neuropixels, pose specific challenges for data preprocessing and the achievement of high-quality spike sorting. The Power Pixel pipeline recapitulates the steps in the pipeline used by the International Brain Laboratory (International Brain Laboratory et al., 2022). The backbone for this is the SpikeInterface implementation of the preprocessing steps, these are all done “lazily”. This means that the result of each step does not result in a new raw binary file which is saved on the disk, instead SpikeInterface creates a virtual processing pipeline of all the steps which is executed and saved to disk only once, just before spike sorting. This preprocessed raw binary file is subsequently deleted after the spike sorting is done to save disk space. This means that these preprocessing steps only impact spike sorting output, the raw data remains unchanged after the pipeline has run.

#### High-pass filter

Raw data that is recorded in broadband, such as is the case with Neuropixel 2.0 probes, first has to be high-pass filtered. To this end, the first step of the pipeline is to apply a 300Hz high-pass Butterworth filter.

#### Inter-sample shift correction

On a Neuropixel probe, the active channels are not sampled exactly simultaneously. Instead, groups of channels are sampled consecutively within each sample acquisition. This results in a shift in the acquisition time between channel groups in the order of tens of microseconds. These shifts, albeit minuscule, can cause artifacts while applying a common-average reference (CAR) over the entire probe. Therefore, these shifts are corrected by the pipeline.

#### Detect and remove channels

Before performing any common average referencing the channels that are noisy, or outside of the brain, should be removed, this prevents these signals from contaminating the reference signal. First, dead channels and channels outside of the brain are detected and removed. Subsequently, common average referencing is used to denoise the remaining channels, channels that are still noisy after CAR are detected and interpolated over neighboring channels (Figure 2, step 1). The method by which bad channels are detected is described in detail in International Brain Laboratory et al., 2022.

**Figure 2.**
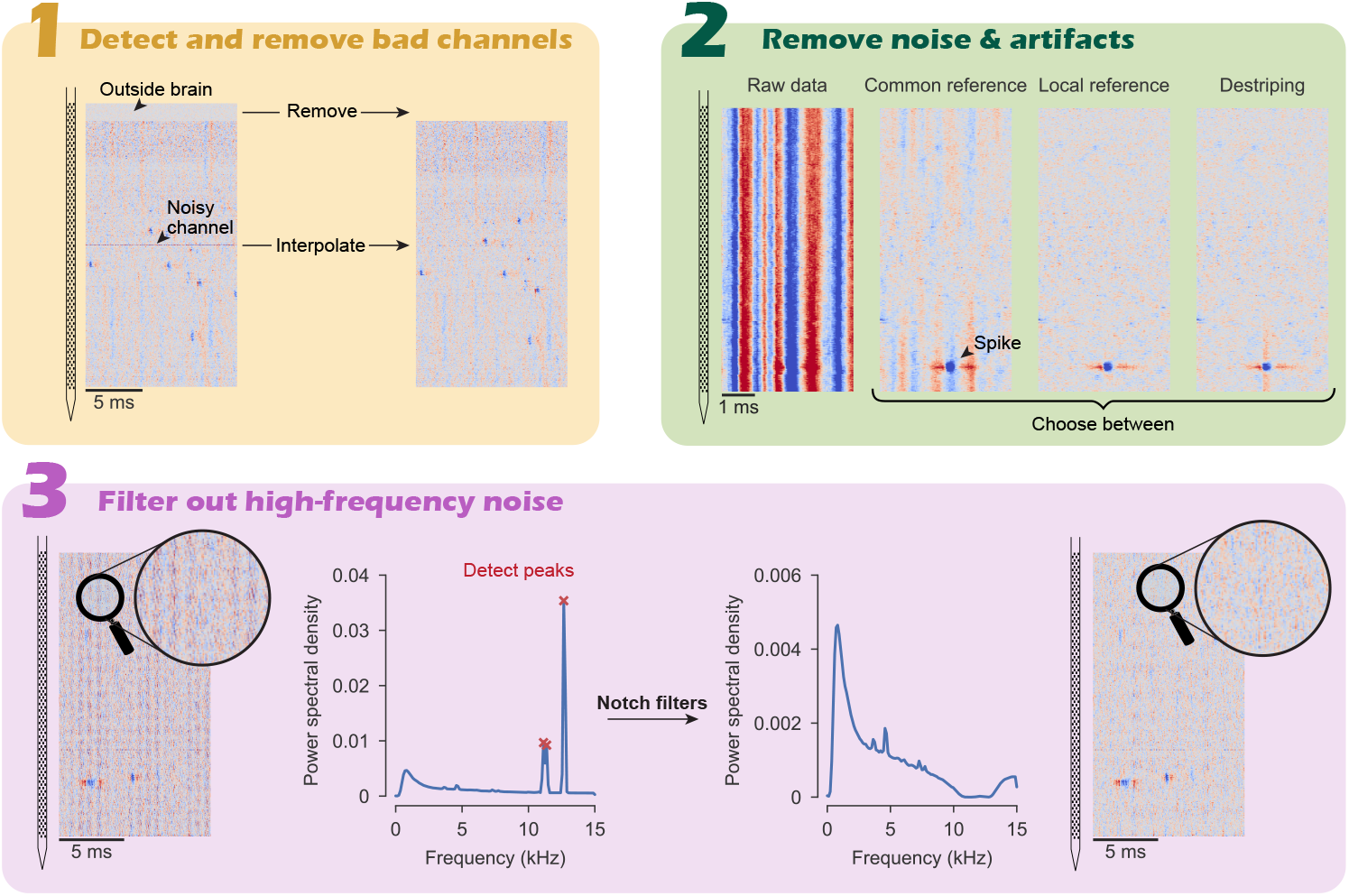
Preprocessing steps the pipeline employs before starting spike sorting

#### Artifact removal

Electrical artifacts can have many different sources, but they share one feature: their signal is the same over nearby channels. Electrical artifacts which are identical across all channels of the probe can be filtered out using common average referencing. Neuropixel probes, however, are so long that electrical artifacts can have slightly different amplitudes at the tip versus at the base of the probe. Therefore, performing a common average reference, whereby the median of all channels is subtracted from each channel, can be insufficient to filter out electrical noise (Figure 2, step 2 *Common reference*). Therefore, the user has the option to employ a spatial filtering method developed by the IBL called *destriping*. This method takes into account that artifacts can vary smoothly over the length of the probe. Besides destriping, a third option is to apply a *local reference* whereby the median of channels in close vicinity of each channel are subtracted from it. Local channels are selected by drawing an annulus around each channel with an inner diameter of 50 *µ*m and an outer diameter of 200 *µ*m (these values can be changed in the settings), channels are selected which fall inside the inner and outer diameter of the annulus. This approach ensures that neighboring channels are not subtracted from each other since they might contain highly similar signals which could cancel each other out. The user can set which artifact removal strategy they wish to employ: common average referencing, destriping, or local average referencing (default).

#### High-frequency noise removal

Recordings sometimes have high frequency noise in specific frequency bands. These are automatically detected in the power spectrum of the recording and removed using notch filters targeted to the peaks in the power spectrum (Figure 2, step 3). A plot of the power spectrum with the detected peaks is saved in the session folder together with a plot of the power spectrum after the notch filters have been applied. Peak detection is done with scipy’s find_peaks, the threshold value of the peak detection can be set by the user if they are not happy with the default threshold.

### Spike sorting

The pipeline is developed using Kilosort (Pachitariu et al., 2024, 2016), other spike sorters will probably work as well but have not been tested. Files with the recommended parameters for Kilosort 2.5, 3, & 4, and pykilosort are included in the pipeline repository but can be changed if necessary. Spike sorting is run by SpikeInterface which allows seamless integration in the pipeline, MATLAB-based sorters (such as Kilosort 2.5 and 3) are deployed in a Docker container, alternatively the user can install Python-based Kilosort 4 locally.

### Neuron-level QC metrics

Modern spike sorters, like Kilosort, tend to be overzealous when detecting units. Only a subset of the detected units are well isolated single units, with the majority of detected units being multiunit activity or noise. Therefore, after spike sorting has detected putative single units, quality metrics are calculated for these units. These metrics give an indication of how clean the unit is, for example, by computing the number of inter-spike interval violations. On the other hand, metrics can also be used to characterize units, for example by looking at the spike width to classify units as putative narrow-spiking interneurons.

### Manual and automatic curation of spike sorting output

To make a final decision on which units constitute single neurons, most electrophysiologists manually curate their data. This entails visual inspection of the waveforms, interspike interval distribution, spike amplitudes, etc. Power Pixels computes many of these quality metrics to provide the user with as much information as possible upon which to make this decision. The curation GUI of SpikeInterface is used to this purpose. Besides manual curation, however, there are now more and more algorithms which aim to automate this process. The pipeline runs three of these algorithms. The predictions of these algorithms are compiled, together with the indication from Kilosort, and added to the [SpikeInterface manual curation GUI](https://github.com/SpikeInterface/spikeinterface-gui) (Figure 3). In the manual curation GUI the user can use all the metrics computed by the pipeline together with the predictions of the three automated curation algorithms to make their decision regarding which units constitute well isolated single neurons. In the GUI this can be indicated with a drop-down menu per neuron, allowing the user to specify each unit as single neuron, MUA, or noise. These choices are saved to disk and the *load_neural_data* helper function has an option to only load in neurons that are deemed single neurons by the user. Alternatively, one can use the predictions of one of these automatic curation algorithms and skip the manual curation step altogether.

**Figure 3.**
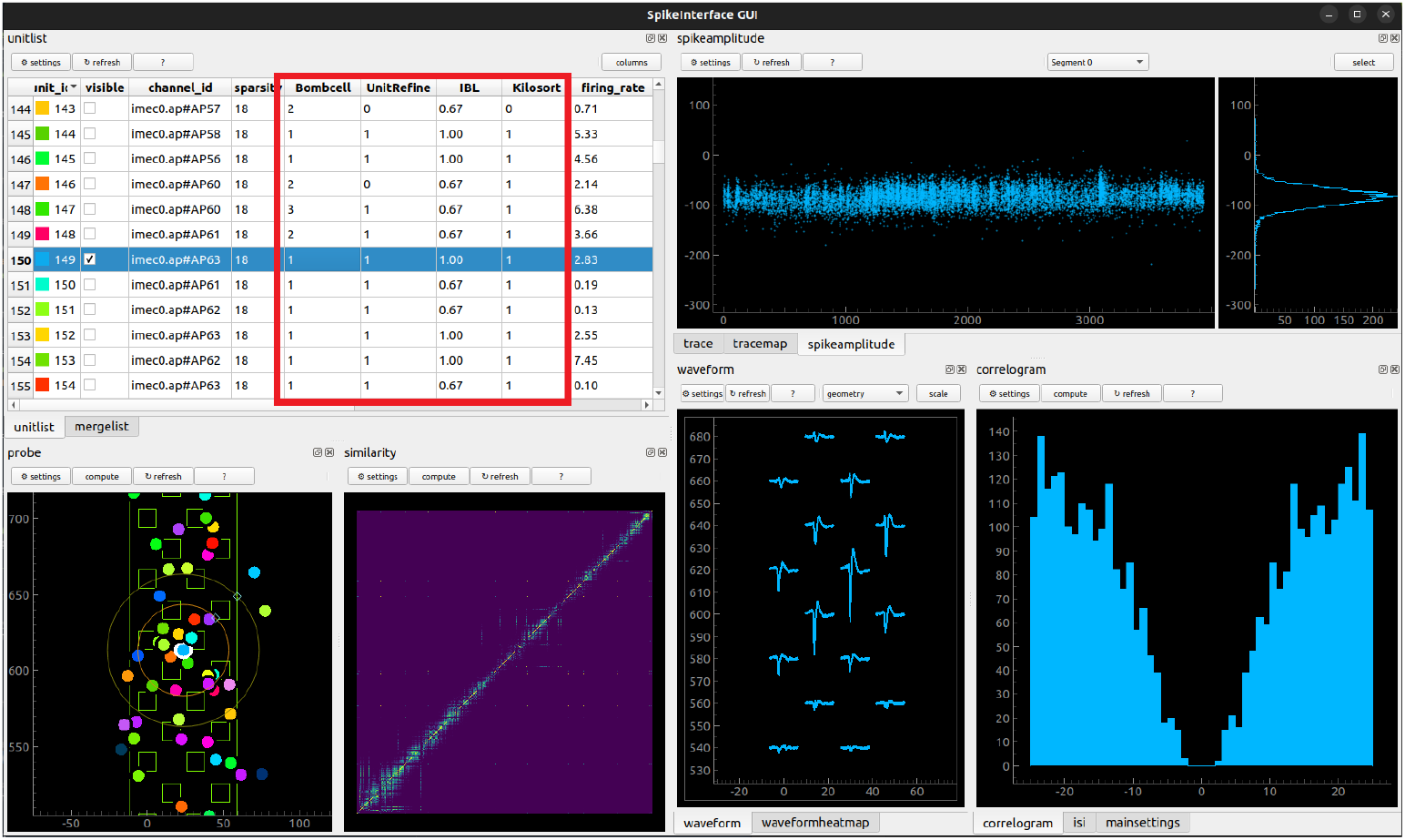
The manual curation GUI of SpikeInterface (developed by Samuel Garcia) with added predictions of which units are well isolated single units from several algorithms (outlined in red).

#### Bombcell

A toolbox which classifies units as good, multi-unit activity, noise, or non-somatic based on a variety of spiking features (Fabre et al., 2023). In short, Bombcell goes through a series of decision steps sequentially to determine unit identity (Figure 4). First, some basic waveform features are checked to ensure the waveform of the unit matches what is to be expected of a real neuron, if it fails one of these checks the unit is classified as noise. Secondly, a check is done to see if the neuron has a large postitive peak which is indicative of an axonal or dendritic spike instead of a spike originating from a soma. Lastly, the remaining units are classified as single neurons or MUA depending on, for example, refractory period violations. All the metrics listed in Figure 4 are exposed to the user in the *bombcell_params*.*json* file which is generated by the pipeline, and can be set according to the needs of the user. The output of the classification needs to be in integers to be ingested by the manual curation GUI, the following mapping is used: 0 = noise, 1 = good single neuron, 2 = multi-unit activity, 3 = non-somatic (i.e. axonal).

**Figure 4.**
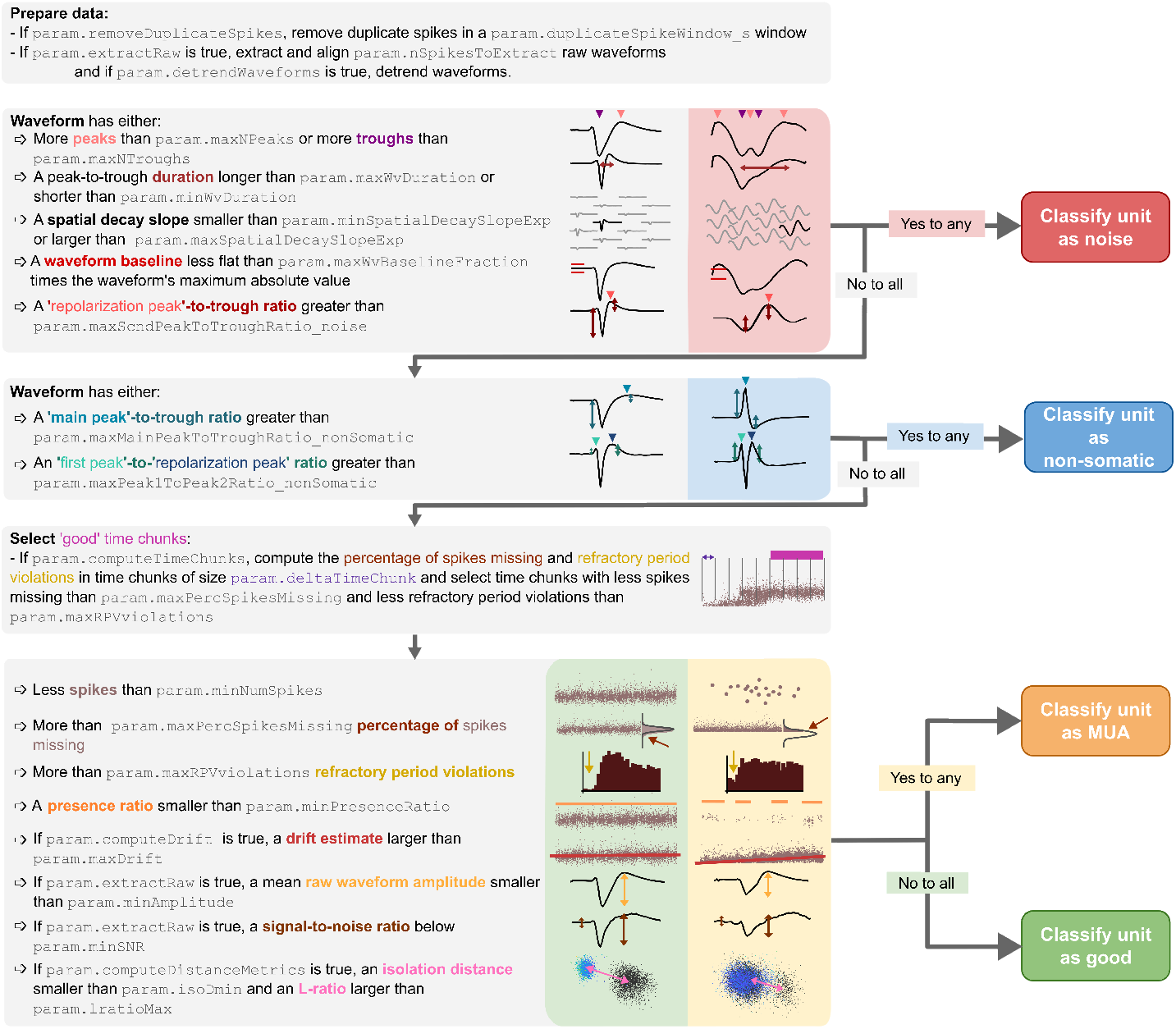
Flowchart depicting when neurons are classified as noise, non-somatic, MUA, or good by Bombcell. From Fabre et al., 2023

#### UnitRefine

This is a Random Forest classifier trained on recording sessions which were manually annotated by two expert humans (Jain et al., 2025). The training data consisted of 11 recordings from visual cortex, suprior colliculus, and motor cortex of the mouse performed with Neuropixel 1.0 probes. The classifier has been pretrained to identify which combination of features are the best predictors of whether a unit is a single neuron. Because this is a pretrained machine learning classifier, it does not have any tunable parameters. The upside of this is that the user does not have to manually tune the parameters to fit their needs, the UnitRefine publication claims that it achieves human-level performance without the need for manual parameter tuning. The downside is that there is no control over the output of the classification which might be an issue for users which value this.

Classification can be performed in two ways: (1) a specialized classifier first classifies noise from neural units, subsequently only the neural units are divided into single units and multi-unit activity by a different classifier, and (2) only the single unit classifier is used on all the units. The default behavior of the pipeline is to apply the latter case in which only single units are classified from the rest, this is because two-stage (or cascaded) classification suffers from compounding errors; the misses and false alarms of the first classification step are propagated to the second classification step resulting in a higher error rate at the end. However, if the user is specifically interested in multi-unit activity, the noise classification can be turned on in the settings. In both cases the output of the classifier is as follows: 0 = noise (if applicable), 1 = single unit, 2 = multi-unit activity / noise.

As mentioned above, the single unit classifier is trained on data from mice. There are, however, pretrained classifiers available which are trained on neural data from rats, naked mole rats, monkeys, and humans. The user can decide to switch from the default classifier (trained on mice data) to any of the other classifiers by changing the used model in the UnitRefine parameters to any of the other available ones (see [here](https://huggingface.co/AnoushkaJain3)).

#### IBL metrics

Neuron-level QC metrics developed by the IBL which evaluate units on three criteria: refractory period violations, amplitude distribution cut-off, and spike amplitude (International Brain Laboratory et al., 2022).

Refractory period violations are typically defined as spikes which occur within a fixed time period, often 2 ms, after another spike. However, refractory period lenghts are known to vary with neuron type and brain region. To overcome this, a range of possible refractory period lenghts (0.5-10 ms, 0.25 ms bins) is used for which the maximum number of acceptable violations is calculated given a chosen contamination level (default is 10%). The confidence that a neuron is less than 10% contaminated must be over 90%, given Poisson spiking, to pass this criterion.

The amplitude distribution cut-off criterion looks for cases in which the distribution of spike amplitudes is “cut off” at the tail, indicating the neuron has spikes that are missed because their amplitudes did not reach the threshold for detection. To this end, the amplitudes in the lowest bin of the amplitude distribution are compared to the amplitudes of the highest bins. To pass this criterion, the lowest amplitudes must be 5 standard deviations lower than the highest amplitudes. Furthermore, the lowest bin must be less than 10% of the highest bin of the distribution.

The median spike amplitudes must be larger than 50 *µ*V to pass this criterion. All the parameters are tunable by the user in the *ibl_qc_params*.*json* file and are summarized in Table 1. If a unit passes all three criteria it gets a score of 1, if it passes two out of three criteria a score of 2/3, one of the three 1/3 and if it passes none of these it gets a score of 0.

**Table 1.**
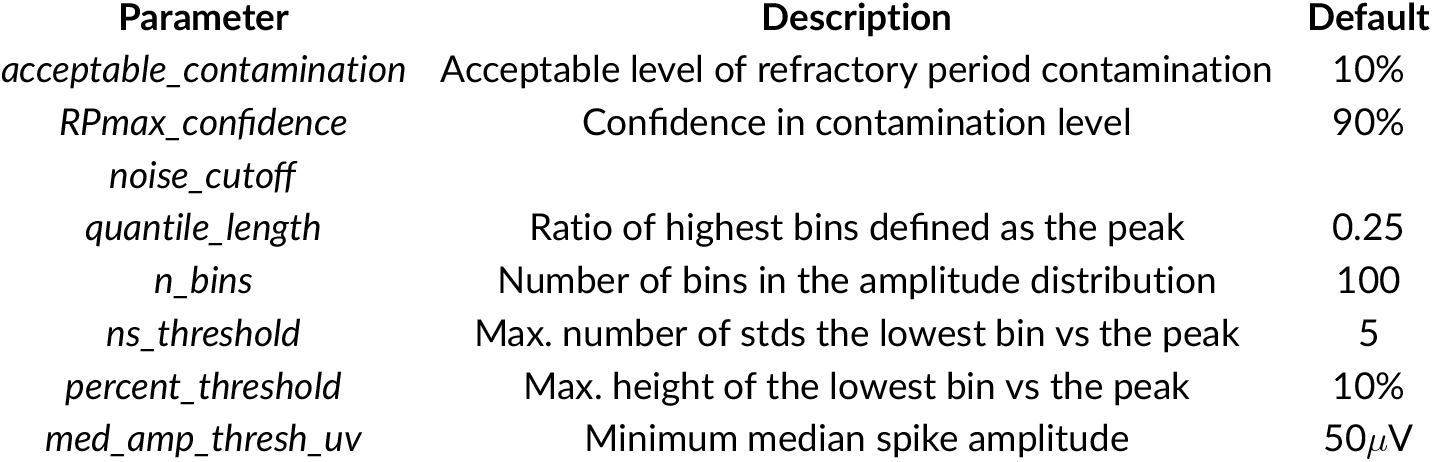
Parameters of the IBL neuron-level QC metrics.

| Parameter | Description | Default |
| --- | --- | --- |
| *acceptable_contamination* | Acceptable level of refractory period contamination | 10% |
| *RPmax_confidence* | Confidence in contamination level | 90% |
| *noise_cutoff* |  |  |
| *quantile_length* | Ratio of highest bins defined as the peak | 0.25 |
| *n_bins* | Number of bins in the amplitude distribution | 100 |
| *ns_threshold* | Max. number of stds the lowest bin vs the peak | 5 |
| *percent_threshold* | Max. height of the lowest bin vs the peak | 10% |
| *med_amp_thresh_uv* | Minimum median spike amplitude | 50μV |

#### Parameter tuning

The two rule-based automatic curation approaches (Bombcell and IBL) have a large number of tunable parameters, as described above. The default parameters are chosen by the respective developers to fit the majority of use cases. However, the default parameters most likely reflect the predominant use case of Neuropixel recordings: recordings in the forebrains of mice. If the user performs recordings in a different species, or a brain region with atypical neural activity, these default parameters might fail. In this case the best course of action is to manually annotate several recordings and run the curation algorithms with different parameters until a good fit between the automatic classification and the manual classifcation is observed. This is also a good approach to determine which of the three automatic curation methods works best on your data. To facilitate reproducibility, the user is encouraged to publish the parameters of the automatic curation method they use in their manuscript.

### Raw electrophysiology metrics

Besides spiking activity, other information can be extracted from the raw electrophysiological traces of each channel. From the low-frequency component of the raw voltage trace, the local field potential (LFP), the power in different frequency bands is computed using spectral decomposition. Furthermore, the root-mean square (RMS) of the high-frequency component is computed.

### Histology

Another important part of the Power Pixels pipeline is to determine which brain region each recording channel is sampled from. To this end, the experimenter should label their probes with a fluorescent dye (typically Di-I [V22888, ThermoFisher Scientific]) before insertion into the brain. After multiple recording penetrations are carried out, the experimenter will typically perfuse the animal, extract the brain, slice the brain, mount the brain slices on slides, and image these slides with a fluorescence microscope. This results in a collection of coronal slices in which the fluorescence tract of the probe is visible. The next step is to map the brain slices to the Allen Brain atlas and trace the fluorescence tract in these registered slices. The pipeline supports two MATLAB packages specifically designed to manually align histological slices to the Allen Brain Atlas and trace probe tracts there: AP_Histology andUniversal Probe Finder (Montijn Heimel, 2022). After the experimenter has done this, the pipeline includes a conversion functions to convert the coordinates of the probe trajectories to a format that can be read by the Ephys-Histology Alignment GUI developed by the IBL.

#### MATLAB requirement

To our knowledge, there is no Python package available that can align coronal brain slices to the Allen Atlas and trace fluorescent probe track through them. The Python-based option used by the IBL called [lasagna](https://github.com/SainsburyWellcomeCentre/lasagna), only works by registering the full 3D brain stack to the Allen Atlas. This requires whole-brain imaging using a serial-sectioning two-photon scanner or a light sheet fluorescence microscope. Both devices are highly expensive and will not be available to the typical experimenter. This means that, for this step of the pipeline only, the user needs a MATLAB licence.

### Ephys-Histology alignment

Probe tracts in the brain can be well defined in terms of their insertion vector using the method described above. However, their precise depth cannot be reliably determined solely on the basis of the fluorescence tract. This is for several reasons: (1) the fluorescent dye diffuses at the tip making it hard to determine where the tip exactly was, (2) the experimenter might pull back the probe slightly before recording to increase recording stability, and (3) the force exerted on the brain by the probe can result in non-homogeneous compression of the brain tissue. Therefore, to establish which recording channel was in which brain region, the brain regions along the insertion vector need to be aligned with the electrophysiological markers recorded by the probe. For example, LFP power is high in the dentate gyrus but low in fiber bundles. Also, cross-correlations between neurons are expected to be higher within brain regions compared to across. To do this alignment, the ephys-histology alignment GUI developed by the IBL will be used (documentation can be found [here](https://github.com/int-brain-lab/iblapps/wiki)). The GUI allows the experimenter to move, stretch and squeeze the brain region until they fit with the recorded electrophysiological markers.

### Synchronization

When using multiple Neuropixel probes and / or when using a [BNC breakout board](https://www.ni.com/nl-nl/support/model.bnc-2110.html) (to record timestamps from behavioral events), the spike times need to be synchronized between the probes and with the event times. This is because each probe, and the BNC breakout board, has its own internal clock. These clocks can show small shifts over time which need to be corrected. To this end, the Neuropixels PXI basestation generates a 1s square wave. This synchronization signal is automatically routed to the headstages of all the probes, the (optional) BNC breakout board should receive the square wave by connecting the SMA connector of the PXI to one of the digital inputs of the breakout board (Figure 5). Alternatively, one can generate a custom synchronization pulse using an Arduino and use that for synchronization. For more details and instructions see the documentation [here](https://billkarsh.github.io/SpikeGLX/help/syncEdges/Sync_edges/) and [here](https://open-ephys.github.io/gui-docs/Tutorials/Data-Synchronization.html). The spike times of multiple probes are synchronized to each other by aligning each probe to the master clock of the PXI basestation. This is done by linearly interpolating the spike times relative to each toggle of the 1s square wave generated by the basestation. Since the square wave signal is recorded by all probes, and the BNC breakout board, it can be used to align all signals. Clock drifts are effectively canceld out by using each toggle of the square wave as an anchor point from which to perform linear interpolation. All this is done under the hood and propagated to the spike times in the final data files.

**Figure 5.**
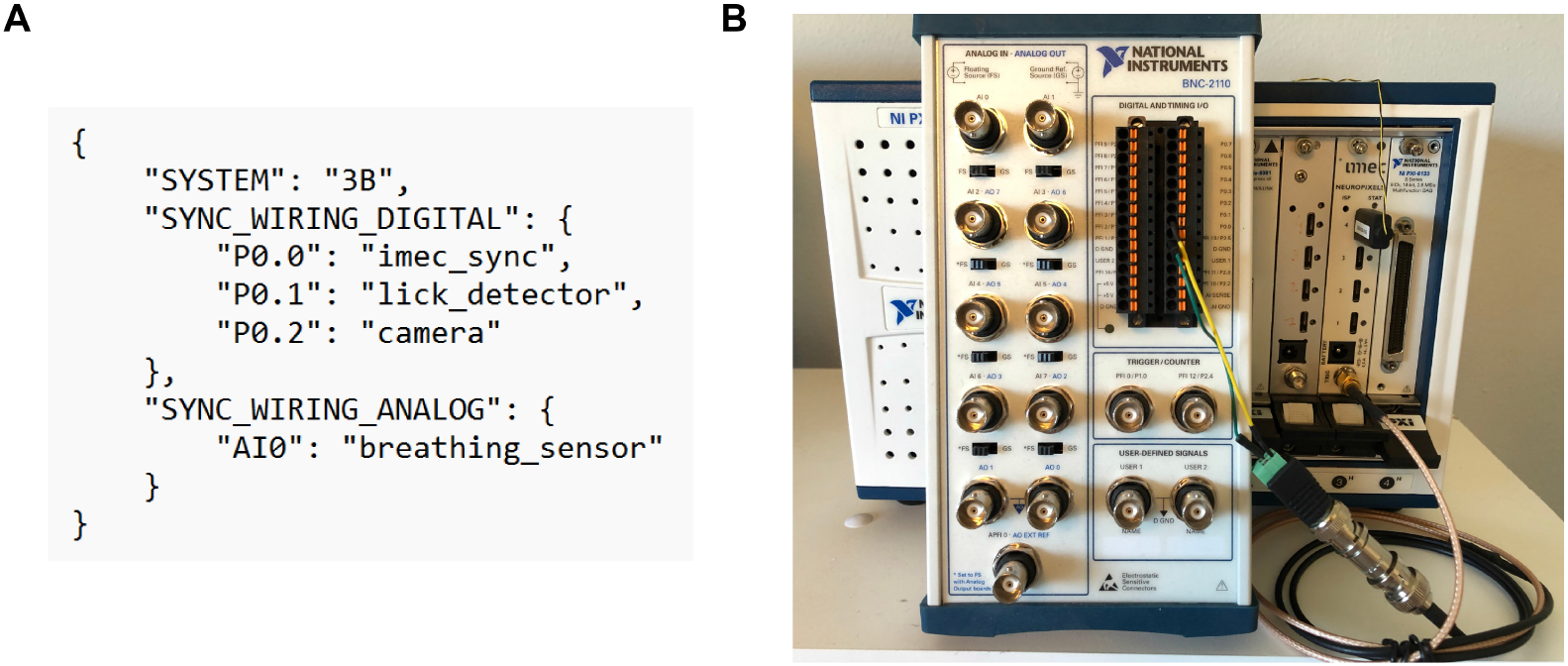
(A) Example JSON file describing the wiring of the BNC breakout box. In this example the 1s square wave pulse from the PXI chassis is relayed to the first digital channel (P0.0). The next two digital channels are used to record timestamps of licks and camera frames. The first analog channel is used to record a continuous signal from a breathing sensor. (B) A picture of the BNC breakout box (BNC-2110, National Instruments) which receives the synchronization pulse from the PXI chassis as an input in the first digital channel (P0.0). Picture credits [Open Ephys](https://open-ephys.github.io/gui-docs/Tutorials/Data-Synchronization.html).

### Data output

The output of the pipeline is in (1) [ALF data standard files](https://int-brain-lab.github.io/iblenv/docs_external/alf_intro.html), (2) NWB format (optional), and (3) native Kilosort files. The ALF data standard is a file naming convention which dictates how files should be called and organized. The files themselves are regular .*npy* files which can be easily loaded in python. For example, the *spikes*.*times*.*npy* file contains an array of all the spike times of all units and the *spikes*.*clusters*.*npy* is an array of equal length containing all the neuron identities. The advantage of this naming convention is that the data can be easily shared using the [Open Neurophysiology Environment (ONE)](https://int-brain-lab.github.io/iblenv/one_docs/one_reference.html). Furthermore, having the data in this format allows one to use all the functions from the Brainbox library, which is part of [ibllib](https://github.com/int-brain-lab/ibllib). These functions include everything from plotting of peri-stimulus time histograms to decoding. For a detailed explanation of all the dataset types found in the output folder refer to [this table](https://docs.google.com/spreadsheets/d/1ieLXRPLLSgUKcLvFkrqizfZl5HjdfE6bQ2KLBCRmjQo/).

## Conclusion

The Power Pixels pipeline provides researchers the same integrated experience, while processing Neuropixel recordings, as those working in large collaborations. It uses SpikeInterface and the IBL codebase to mimic the tried-and-tested pipeline used by the International Brain Laboratory. This pipeline is especially useful for researchers who are new to Neuropixel recordings and might not have extensive insight into which choices to make during the processing of these recordings. After initial setup has been done the pipeline runs almost completely automatically through all the preprocessing steps and the spike sorting. The only preprocessing step which requires manual intervention is determining whether high frequency noise is present in the recording which needs to be filtered out with targeted notch filters. The biggest manual effort is tracing the probe tracts through the histological brain slices and subsequently aligning electrophysiological features to the brain regions determined by histology. To conclude, the Power Pixels pipeline requires minimal human intervention to go from raw data to spike sorted single neurons in specified brain regions.

## Code availability

The Power Pixels pipeline code is available on Github:

https://github.com/NeuroNetMem/PowerPixelsPipeline/

## Acknowledgments

We thank all the developers and contributors to the following open-source projects: SpikeInterface (special thanks to Alessio Buccino and Samuel Garcia), The International Brain Laboratory (special thanks to Oliver Winter and Mayo Faulkner), Bombcell (Julie Fabre), UnitRefine (Anouska Jain), Universal Probe Finder (Jorrit Montijn), AP_Histology (Andy Peters), and Kilosort. This work was supported by the Dutch Research Council (NWO) through grant number OCENW.XL21.XL21.069.

